# Evolution of alternative reproductive systems in *Bacillus* stick insects

**DOI:** 10.1101/2023.07.31.550487

**Authors:** Guillaume Lavanchy, Alexander Brandt, Marc Bastardot, Zoe Dumas, Marjorie Labedan, Morgane Massy, William Toubiana, Patrick TranVan, Andrea Luchetti, Valerio Scali, Barbara Mantovani, Tanja Schwander

## Abstract

Reproduction is a key feature of all organisms, yet the way in which it is achieved varies greatly across the tree of life. One striking example of this variation is the stick insect genus *Bacillus*, in which five different reproductive modes have been described: sex, facultative and obligate parthenogenesis, and two highly unusual reproductive modes: hybridogenesis and androgenesis. Under hybridogenesis, the entire genome from the paternal species is eliminated, and replaced each generation by mating with the corresponding species. Under androgenesis, an egg is fertilized but the developing diploid offspring bear two paternal genomes, and no maternal genome, as a consequence of unknown mechanisms. Here, we re-evaluate previous descriptions of *Bacillus* lineages and the proposed F_1_ hybrid ancestries of the hybridogenetic and obligately parthenogenetic lineages (based on allozymes and karyotypes) from Sicily, where all these reproductive modes are found. We generate a chromosome-level genome assembly for a facultative parthenogenetic species (*B. rossius*) and combine extensive field sampling with RADseq and mtDNA data. We identify and genetically corroborate all previously described species and confirm the ancestry of hybrid lineages. All hybrid lineages have fully retained their F1 hybrid constitution throughout the genome, indicating that the elimination of the paternal genome in hybridogens is always complete and that obligate parthenogenesis in *Bacillus* hybrid species is not associated with an erosion of heterozygosity as known in other hybrid asexuals. Our results provide a stepping stone towards understanding the transitions between reproductive modes and the proximate mechanisms of genome elimination.

## Introduction

The most widespread form of reproduction in animals is sexual reproduction with separate male and female sexes. Sexual reproduction involves the formation of gametes via meiosis, which fuse with gametes from another individual to form a zygote (syngamy). However, modifications of the two key processes within meiosis, recombination and segregation, as well as partial or complete suppression of syngamy, have generated alternative reproductive strategies, including parthenogenesis, hybridogenesis and androgenesis (Figure 1). The most frequent alternative reproductive strategy is parthenogenesis (which can be obligate or facultative), where females produce offspring from unfertilized eggs. It is known from natural populations of all major animal lineages except mammals. In addition to the absence of fertilization, parthenogenesis involves different modifications of meiosis, which vary among species (Suomalainen et al., 1987; Neiman et al., 2014). In contrast to parthenogenesis, hybridogenesis (Lavanchy & Schwander, 2019; Schultz, 1969) involves the production of offspring from fertilized gametes. Recombination between parental chromosomes does not occur and only the chromosomes inherited from one of the parents are transmitted to the gametes. The excluded parental genome is then replaced at every generation via mating with a sexual species. Hence, hybridogenetic lineages need to co-occur with populations of a sexual species in order to have access to mating partners. Hybridogenesis is known in a few insect lineages (Hamilton et al., 2018; Mantovani & Scali, 1992), as well as in different fish and frog taxa, where it has been studied most extensively (Schultz, 1969; Uzzell & Berger, 1975; Schmidt et al., 2011; Kimura-Kawaguchi et al., 2014). Finally, under androgenesis, offspring bear only nuclear genes from their father, as there is no fusion between maternal and paternal pronuclei and only paternal pronuclei contribute to the zygote. In haplodiploid species, where androgenesis is the most widespread, only males (which are haploid) have been found to be produced by androgenesis. In other taxa, male, female, or hermaphrodite offspring can be produced via androgenesis, with two nuclear genome copies stemming from the father (reviewed in Schwander & Oldroyd, 2016).

**Figure 1:**
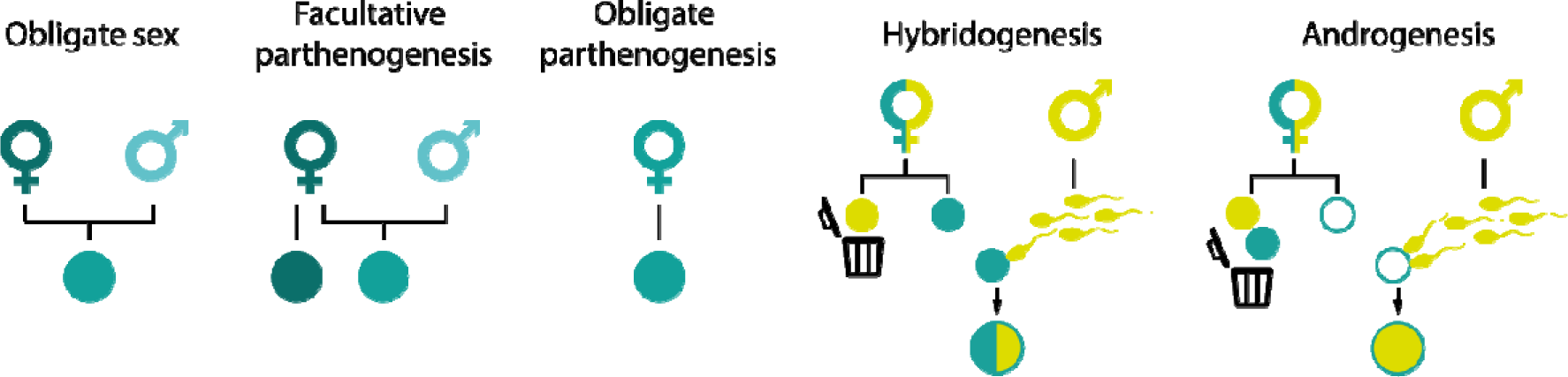
Summary of the different reproductive modes described in the genus *Bacillus*. Different shades of the same color designate different genotypes of the same species, while different colors represent genomes from different species.

A striking example of this diversity in animal reproductive modes is the stick insect genus *Bacillus* (Phasmatodea). The genus comprises different but morphologically similar species and lineages that reproduce via obligate sex, facultative and obligate parthenogenesis, as well as hybridogenesis and androgenesis (Scali and Mantovani, 1989; Mantovani & Scali, 1992; Figure 1). Nine lineages co-occur in Sicily, the largest island in the Mediterranean Sea (reviewed in Scali 2009). Three of these belong to an obligate sexual species, *B. grandii* (subspecies *B. g. grandii*, *B. g. benazzii*, *B. g. maretimi*). A fourth lineage, *B. rossius redtenbacheri*, is facultatively parthenogenetic: females reproduce sexually when mated, but reproduce parthenogenetically in the absence of males (Figure 2A). Three additional lineages are obligately parthenogenetic, *B. atticus*, *B. whitei* and the triploid *B. lynceorum*. Finally, two lineages reproduce via hybridogenesis, as well as occasional androgenesis (Mantovani & Scali, 1992; Tinti & Scali, 1995). Previous work, based on extensive karyotype surveys and allozyme studies, indicated that two parthenogenetic lineages, *B. whitei* and *B. lynceorum*, as well as the two hybridogenetic lineages, might be of hybrid origin between *B. r. redtenbacheri* (maternal ancestor) and *B. g. grandii* (Nascetti and Bullini, 1980; Mantovani et al., 1992). The two hybridogenetic lineages bear chromosome sets of different subspecies of *B. grandii*: *B. g. grandii* (hereafter hybridogen 1; Mantovani & Scali, 1992) and *B. g. benazzii* (hereafter hybridogen 2; Mantovani et al., 1991; Figure 2A, 2B). Whether the different hybrid lineages originated from multiple independent hybridization events or whether they are the product of a single hybridization, which then diversified into several lineages, is unknown. The origin of one of the two hybrid parthenogens *(B. lynceorum*), is believed to involve a second hybridization event, as it is triploid and also carries a chromosome set of *B. atticus* in addition to those from *B. r. redtenbacheri* and *B. grandii* (Mantovani et al., 1992; Figure 2B). However, no genetic study including all lineages and more than a few allozyme markers has been performed thus far.

Here, we revisit the *Bacillus* species and lineages from Sicily, using genome-wide SNP genotyping (RADseq) and a mitochondrial marker, to corroborate the original species descriptions based on karyotype and allozyme data, and characterize the genomic composition of hybrid species and lineages with alternative reproductive modes. Specifically, we test whether these lineages are characterized by a genome-wide interspecific F_1_ hybrid ancestry, or whether different genomic regions feature different levels of introgression from the presumed maternal and paternal ancestors. We generate a chromosome-level genome assembly of *B. r. redtenbacheri*, the presumed maternal ancestor of all hybrid lineages from Sicily. We then genotype 444 field-collected individuals from Sicily, identify the number of different species and lineages present in Sicily and analyze hybrid ancestries along each chromosome.

**Figure 2:**
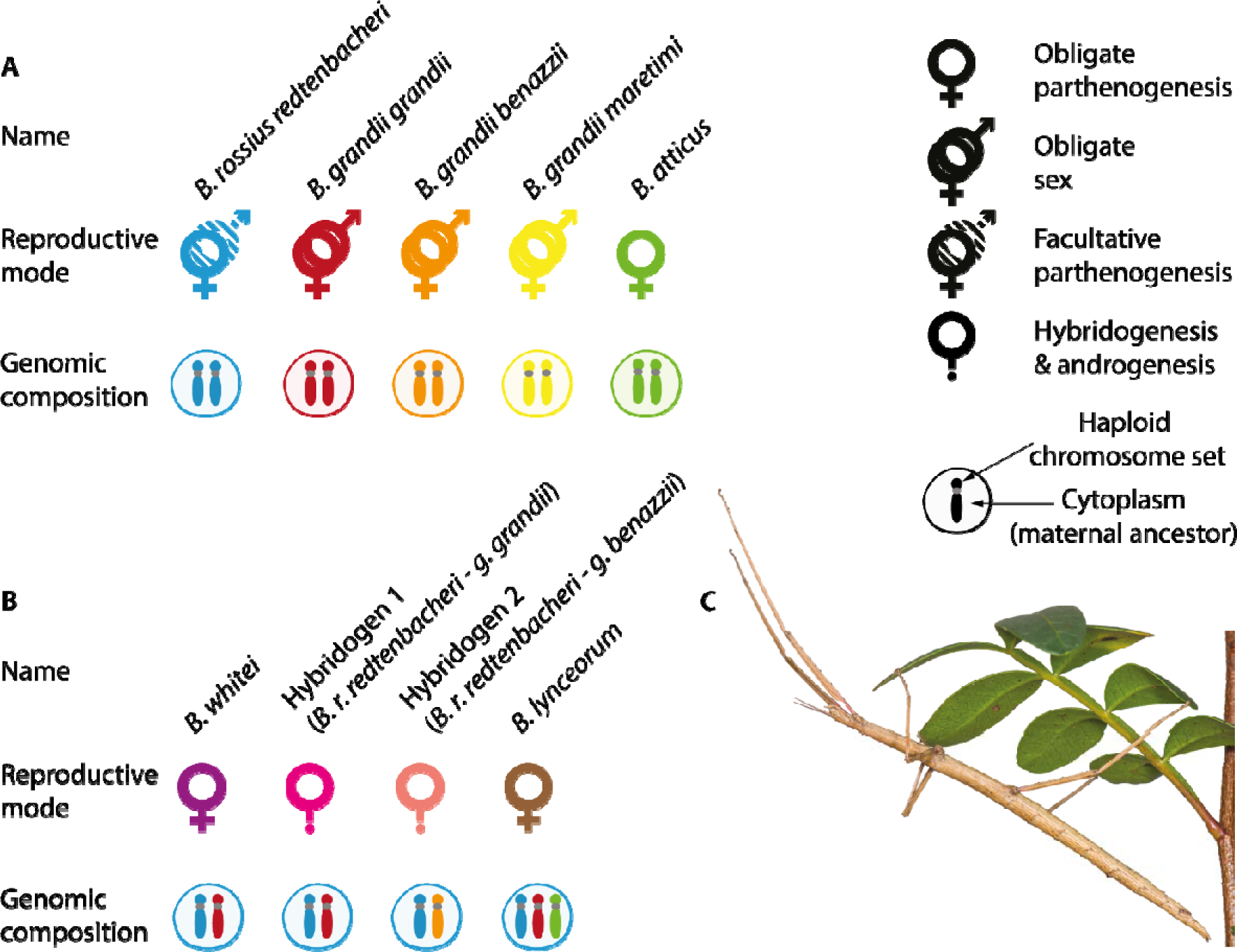
Described reproductive modes and genomic composition of different lineages in the genus *Bacillus*. **A** Reproductive modes and genomic composition of the non-hybrid lineages in the Sicilian area. **B** Reproductive modes and suggested genomic composition of the hybrid lineages, based on allozymes and karyotypes. **C** Picture of an adult *Bacillus* female on one of its host plants, the lentisk *Pistacia lentiscus*.

## Methods

### Sample collection

We collected 444 *Bacillus* stick insects in Sicily in October 2017 and 2018 and September 2020 by searching on bramble (*Rubus* sp.) and lentisk (*Pistacia lentiscus*) bushes at night. We focused mostly on the two areas where hybridogenetic individuals were described (Scali et al., 1995). Nine individuals died before we could sequence them and we sequenced their offspring instead (we refer to them as field-collected from here). We collected additional female *B. r. redtenbacheri* from Ravenna, mainland Italy. We used one for reference genome assembly and included seven to our RADseq dataset to assess the genetic distance between the Ravenna and Sicilian populations. The exact sampling location and year of all genotyped individuals are given in Table S1.

### Reference genome assembly

We built a chromosome-level genome assembly for *B. r. redtenbacheri* using Hi-Fi long read sequencing and Hi-C scaffolding. To extract high molecular weight (HMW) DNA we flash-froze the posterior half of the female from Ravenna (without gut) in liquid nitrogen and ground it using a Cryomill (Retsch). We then extracted HMW DNA using a G/20 Genomic Tips kit (Qiagen) following manufacturer’s protocols. We checked DNA integrity on a pulse field agarose gel. Hi-Fi library preparation and sequencing were done at the Lausanne Genomic Technologies Facility (Lausanne, Switzerland) using a SMRTbell Express Template Prep Kit 2.0 and two SMRT cells on the Sequel II (Pacific Biosciences). Hi-C library construction using the Proximo Hi-C Kit and sequencing (250 Mio read pairs) was outsourced to Phase Genomics (Seattle), with ground tissue from the anterior half of the same individual used for cross-linking following the manufacturer’s protocol.

We assembled PacBio Hi-Fi reads into contigs using *Hifiasm* v0.14.1 (Cheng et al., 2021) with default parameters and decontaminated it using *BlobTools* v1.0 (Laetsch & Blaxter, 2017) under the taxrule “bestsumorder”. We generated a hit file after a *blastn* v2.10.1+ against the NBCI nt database, searching for hits with an e-value below 1e-25 (parameters: *-max_target_seqs 10 -max_hsps 1 -evalue 1e-25*), and removed contigs without hits to metazoans from the assembly.

We then mapped Hi-C reads against the decontaminated assembly using *Juicer* v1.6 (Durand et al., 2016) with the restriction site DpnII. Finally, we used *3D-DNA* v180922 (Dudchenko et al., 2017) for chromosome-level scaffolding with parameters *-i 5000 --editor-repeat-coverage 5*. We assessed the completeness of the genome assembly with BUSCO v5.1.2 (Seppey et al., 2019) against the *insecta_odb10* dataset using the *--long* and *--augustus* parameters.

### COII sequencing and SNP genotyping (RADseq)

We extracted DNA from legs (sliced longitudinally) using the Qiagen Biosprint 96 workstation, following the manufacturer’s protocol. We amplified a fragment of the COII mitochondrial gene using the primers 5’-ATGGCAGATTAGTGCAATGG-3’ and 5’-GTTTAAGAGACCAGTACTTG-3’ (Simon et al., 1994). We used 2 µL of DNA extract in 20 µL of solution containing 0.5 µM of each primer, 0.2 mM of each dNTP, 1 U of Q5 HotStart Taq polymerase (New England Biolabs) in 1 X Q5 buffer. We ran the thermocycler for an initial denaturation of 30 s at 98 °C, followed by 30 cycles of denaturation at 98 °C for 10 s, annealing at 51 °C for 30, extension at 72 °C for 30 s, and a final extension at 72 °C for 2 min. The resulting PCR products were checked on gel and sent for Sanger sequencing (forward) at Microsynth (Balgach, Switzerland).

We inspected the sequences and trimmed low-quality extremities in Geneious Prime® 2022.0.2. We added one sequence of each haplotype from Mantovani et al. (2001) as references for taxonomic assignment, including one from *Clonopsis gallica* as an outgroup. We aligned the sequences using MUSCLE3.8.1551. We reconstructed a ML phylogeny with IQ-TREE 2.0.6 (Minh et al., 2020) using ModelFinder Plus (Kalyaanamoorthy et al., 2017) and assessed its robustness with 1000 ultrafast bootstraps.

We prepared double-digest RADseq libraries using a protocol adapted from Parchman et al. (2012) and Brelsford et al. (2016). We digested genomic DNA with restriction enzymes EcoRI and MseI (New England Biolabs) and ligated Illumina primer and adapter sequences to the cut sites with an individual 8-nucleotide barcode on the EcoRI side. We amplified the resulting library in four replicate PCR of 20 cycles. We then size-selected fragments of 300 – 400 bp on a 2.5 % agarose gel. The final library was single-end sequenced (125 bp) on eight Illumina HiSeq 2500 lanes at the Lausanne Genomic Technologies Facility. Individuals collected in different years were included in different libraries, which were prepared at different times.

We first demultiplexed the reads using the *process_radtags* command of stacks v2.3e (Catchen et al., 2013) with the options *-c -q -r -t 143.* We removed remnants of adapter sequences and trimmed bases from 3’ read ends using *cutadapt* v3.4 and *trimgalore* v0.6.6 with default parameters (Krueger, 2021; Martin, 2011). We mapped trimmed reads to the reference genome assembly using *bwa mem* v0.7.17 (H. Li & Durbin, 2010) with the option *-M*. We removed unmapped reads and secondary and supplementary alignments as well as alignments with mapping quality below 20 and sorted the alignments by coordinate using *samtools* v1.12 (H. Li et al., 2009). We called variants using *gatk HaplotypeCaller* (McKenna et al., 2010) with default parameters. Next, we filtered variants using *bcftools* v1.12 and *vcftools* v0.1.14 (Danecek et al., 2011; H. Li, 2011). We retained only biallelic and triallelic SNPs with a quality ≥ 20, and genotypes with a minimum depth of 8. We removed SNPs with a minor allele count < 3 and with > 50% of missing genotypes. Further, we removed 55 individuals with ≥ 75% of genotypes missing. Finally, we removed SNPs with a mean depth > 39.98 (median + standard deviation of per SNP mean coverage distribution) in order to exclude SNPs derived from paralogs that were possibly merged during assembly. The final dataset contained 145’341 SNPs in 396 individuals.

### Genetic identification of species with different reproductive modes and population genetic analyses

Because purely morphological criteria cannot be used to reliably distinguish all *Bacillus* species and lineages, we used a combination of mitochondrial and nuclear data to assign each individual to a species (see Results). We built a phylogeny of the COII gene for all individuals, including references from Mantovani et al. (2001), and defined major nuclear clusters by means of Multi-Dimensional Scaling (MDS) on the RADseq data, using the *isoMDS* function from the R package *MASS* (Venables & Ripley, 2002).

Since the parthenogenetic *B. whitei* and the hybridogen 1 share the same parental ancestors (Figure 2B), we used lab reared individuals with known reproductive mode as references to genetically distinguish field collected individuals. To assess the reproductive phenotype of these reference individuals, we reared them in the absence of males to collect and genotype unfertilized eggs. Unfertilized eggs of hybridogenetic females carry solely one haploset derived from the maternal species *B. r. redtenbacheri,* while unfertilized eggs of parthenogenetic *B. whitei* carry both *B. r. redtenbacheri* and *B. g. grandii* haplosets. We extracted DNA from these females and from batches of one to ten of their eggs. We crushed each egg batch with sterile pipet tips and extracted DNA using the Qiagen BioSprint 96 DNA workstation (Qiagen), following manufacturer protocol for tissue and constructed RADseq libraries as described above. We sequenced the libraries with low coverage on two MiSeq lanes (150bp). We then called SNPs as described above, using these new reference individuals as well as all individuals belonging to *B. g. grandii*, *B. r. redtenbacheri* and their hybrids, filtering for biallelic loci and a minor allele count of 3. We first assessed the ancestry of the unfertilized eggs laid by hybrid reference females using admixture with k = 2, using supervised mode with pure individuals as references. Among these hybrid females, those laying unfertilized eggs of pure *B. rossius* ancestry were considered hybridogens and those laying unfertilized eggs of approximately equal *B. rossius*: *B. grandii* ancestry were considered *B. whitei*. We then assigned our field collected individuals to each of the two lineages based on genetic similarity with our reference individuals. For this, we performed a MDS analysis with all field-collected hybrids (*B. whitei* and hybridogen 1) along with the reference individuals.

### Fine-scale genomic composition of hybrids

We then estimated the fine-scale genomic ancestry of hybrids using *admixture* (Alexander et al., 2009). Because our dataset includes multiple species, some RAD loci are expected to be species-specific, that is, have genotype information for some species but not others. To avoid biasing the genomic composition of hybrids towards the species with the largest number of genotyped loci, we only used loci with genotype information for all three non-hybrid species (*B. grandii*, *B. atticus*, and *B. r. redtenbacheri*). To filter our dataset accordingly, we retained only loci that were genotyped in at least 3 individuals from each of *B. r. redtenbacheri*, *B. g. benazzii*, *B. g. grandii* and *B. atticus*. We excluded the sex chromosome to avoid biasing ancestry estimations in males (see below). We then ran admixture in supervised mode, using pure individuals identified on the MDS as references, in ten replicates, and chose the best replicate based on cross validation error.

We then investigated whether some chromosomes were replaced in hybridogens (e.g., via accidental elimination of a chromosome of *B. r. redtenbacheri* origin) or showed some signs of recombination. For this, we assessed the ancestry of each individual chromosome in each hybrid. For each hybrid lineage, we retained loci that were genotyped in at least three individuals of each subspecies that contributed to this hybrid. We then ran admixture in supervised mode for each chromosome separately, using all individuals from the subspecies that contributed to the hybrid as references.

To assess whether the sex chromosome showed different patterns than the rest of the genome, we used our RADseq data to identify the X chromosome in our genome assembly. For this, we used males and females of the sexual species *B. r. redtenbacheri*, and the two subspecies of the sexual species *B. grandii* with at least ten individuals of each sex. We compared depth between males and females because *Bacillus* have XX/X0 sex determination (Marescalchi & Scali, 1990) and males are expected to show half of the female coverage at the X chromosome. We then used depth ratio at the identified sex chromosomes vs autosomes to sex individuals for which phenotypic sex was not available (young nymphs are difficult to sex; see Supplementary material).

We then looked for local losses of hybrid ancestry at a finer scale in the diploid hybrids. We thus compared observed and expected relative heterozygosity at the genome level. We measured observed relative heterozygosity as the proportion of SNPs genotyped in each individual that were heterozygous. We then estimated expected heterozygosity in F_1_ hybrid individuals by drawing alleles at random from the parental species’ allele frequencies, for 1000 replicates. We did not do it for the triploid *B. lynceorum*, as their triploid constitution would result in higher allelic dropout (because one allele has half the coverage of the other one), which would bias our analysis towards lower heterozygosity. Higher coverage would be needed for these individuals, which we could not obtain since we did not have reliable species identification before analyzing the data.

We then asked whether differences between observed and expected heterozygosity values are restricted to specific genomic locations. For this, we plotted average heterozygosity in 1Mb sliding windows (sliding 500 kb), in each hybrid, together with the average expected value across all replicates.

Finally, we compared relative heterozygosity on the autosomes of the different sexual and parthenogenetic lineages. We split *B. r. redtenbacheri* in sexual and asexual populations, based on the presence of phenotypic males among field-caught individuals. We plotted heterozygosity values using the ridgeline function of the R package *ridgeline* (https://github.com/R-CoderDotCom/ridgeline) and compared them using a Tukey test using the *tukeyHSD* function in R.

## Results

### Bacillus rossius redtenbacheri reference genome

The combination of Hi-Fi sequencing and Hi-C scaffolding resulted in a high-quality, chromosome-level reference genome of *B. r. redtenbacheri,* with 97% of the 1.6 Gbp genome anchored to one of 18 scaffolds (total contig number: 1257, L50=5, BUSCO: Complete:99.1% [Single-copy:98.4%, Duplicated:0.7%], Fragmented:0.7%, Missing:0.2%). Eighteen scaffolds were expected given the haploid genome consists of 18 chromosomes (Scali & Mantovani, 1989). By comparing male / female depth ratio at RADseq loci across the 18 largest scaffolds for the field-collected *B. r. redtenbacheri*, *B. g. grandii* and *B. g. benazzii* (following genetic identification, see below), we inferred that scaffold 2 was the X chromosome in all three lineages (Supplementary information, Figure S1). Looking at depth ratio on the X chromosome vs autosomes enabled us to sex all individuals unambiguously (Figure S2).

### Identification of Sicilian lineages

The combination of the nuclear RADseq data and partial sequencing of the *COII* gene enabled us to assign 393 of the 396 genotyped individuals to all described Sicilian *Bacillus* lineages and species. The three remaining individuals showed intermediate genotypes (see below). The MDS analysis of nuclear RAD loci revealed the presence of eight genetic clusters (Figure 3 A,B). The phylogenetic tree of the partial *COII* gene enabled us to assign seven of the eight clusters to species using published sequences as references (Figure S3, Figure 3F). The remaining cluster comprised the two lineages, *B. whitei* and hybridogen 1, which share a similar genomic composition (see Figure 2B). We were able to unambiguously split this cluster into these two lineages by using reference individuals of each lineage which were identified based on their reproductive mode (parthenogenesis or hybridogenesis; Figure 3C; see Methods for details). Finally, 16 out of 71 *B. g. grandii* individuals carried a mitochondrial haplotype corresponding to *B. r. redtenbacheri* (clustering with *B. g. grandii* in the MDS but with *B. r. redtenbacheri* in the mtDNA tree), which may indicate an androgenetic origin. No individual of putative androgenetic origin was found in *B. g. benazzii*. The number and sex of individuals of each species are summarized in Table 1. Note that our analyses confirmed the ancestry of all the hybrid lineages with alternative reproductive modes as originally suggested from karyotype analyses and allozyme markers (Figure 3E). Furthermore, the MDS did not indicate any notable genetic differentiation between *B. r. redtenbacheri* individuals from Sicily and Ravenna.

**Table 1:**
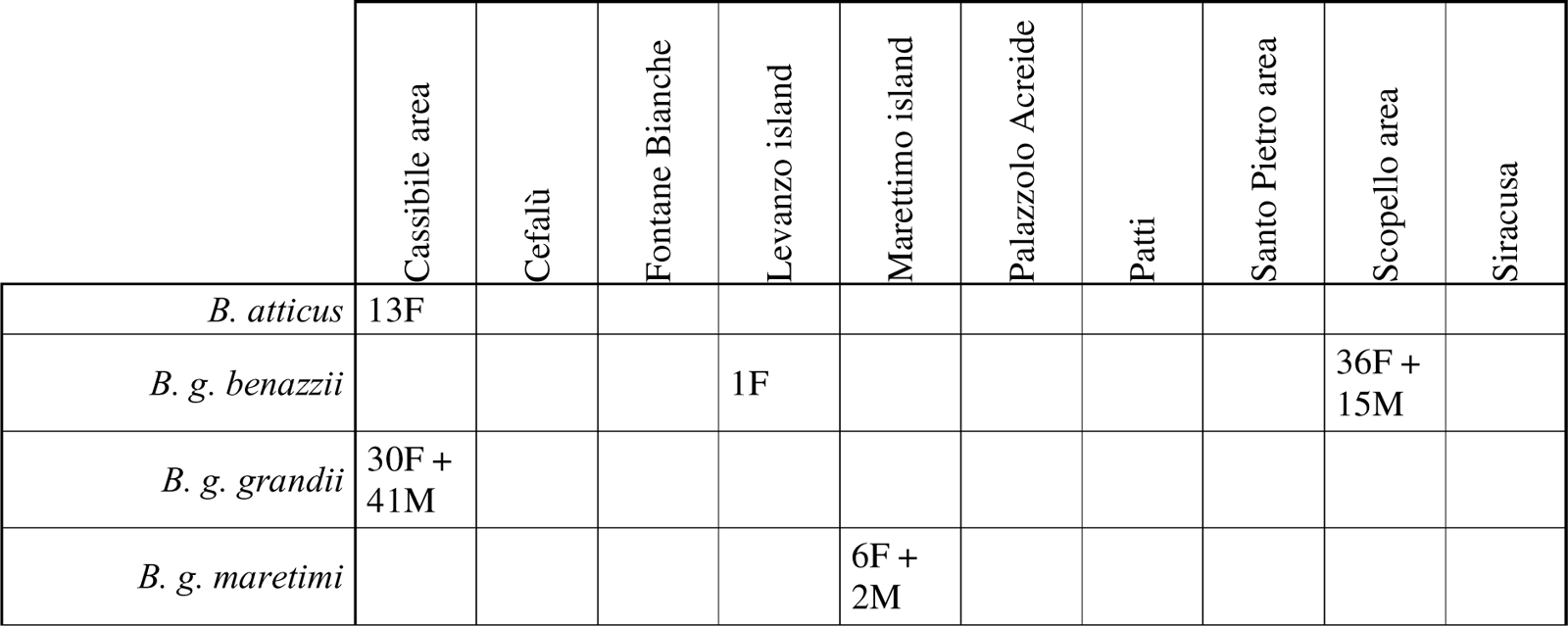
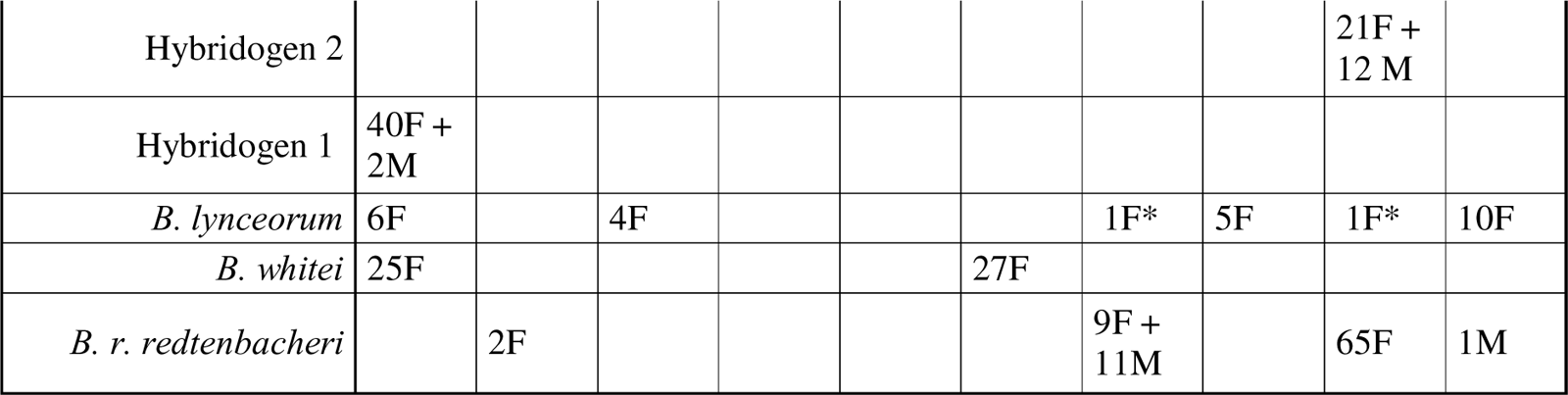
Summary of the number of individuals of each sex collected at each location. F = female; M = male. Sex of juvenile individuals was determined genetically based on the depth ratio between the X chromosome and autosomes (see Supplementary Material, Figure S2). Note that the three novel hybrids (a,b and c in Figure 3) are not included. *Individuals found outside the expected distribution of their species.

**Figure 3:**
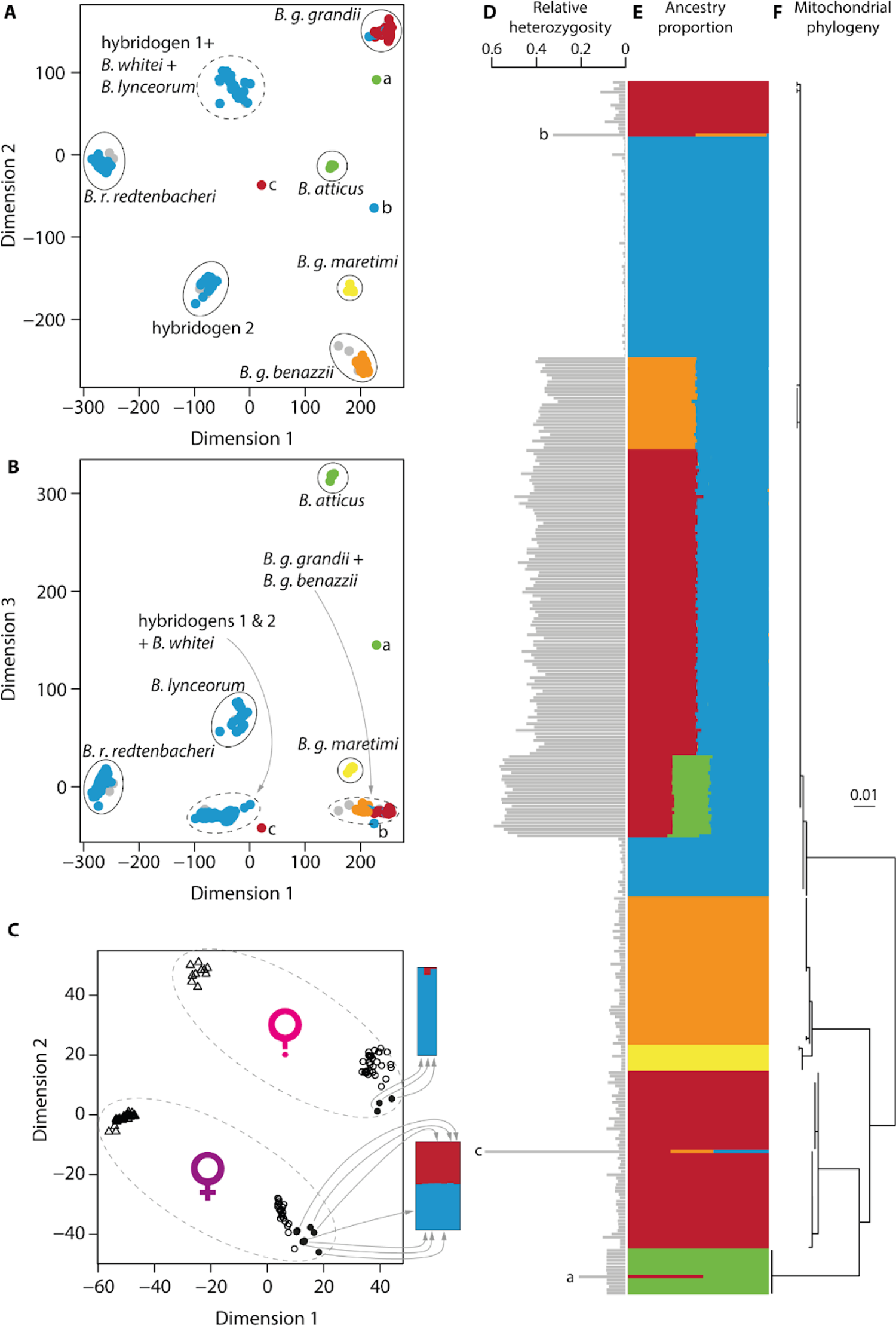
Identification of Sicilian *Bacillus* lineages. **A**: MDS plot of 396 wild-collected individuals, dimensions 1 and 2; **B**: MDS plot of the same individuals, dimensions 1 and 3. Points are colored according to their mitochondrial lineage (see Figure S3). Dimension 1 reflects genetic distance between *B. r. redtenbacheri* and *B. grandi*i/*B. attticus*, dimension 2 between the *B. grandii* subspecies and dimension 3 between *B. grandii* and *B. atticus*. Color code as in Figure 2; grey = COII amplification failed. Clusters are delimited by circles (dashed lines = clusters containing several lineages). a, b and c designate the three individuals falling outside the eight major clusters. **C**: MDS plot of parthenogenetic *B. whitei* and hybridogens 1, including lab-reared individuals for which the reproductive mode was inferred by genotyping their unfertilized eggs. The barplots on the right represent ancestry estimated by admixture for the unfertilized egg batches laid by each female. The correspondence between female and egg batch is shown by arrows. Empty triangles = individuals collected in 2018; empty circles = individuals collected in 2020. Solid circles = lab-reared individuals. **D**: Relative heterozygosity. **E**: Ancestry proportion given by *admixture*; **F**: Maximum-Likelihood phylogeny of 695 bp of the mitochondrial gene COII.

Three of the 396 genotyped individuals, all females, did not belong to any of the eight major clusters of previously described Sicilian lineages but featured intermediate genotypes as expected for different types of hybrids. The first one (a on Figure 3) had the diploid genomic composition of an F_1_ hybrid between *B. g. grandii* and *B. atticus*, with *B. atticus* mitochondria (Figure S7). The second one (b on Figure 3) appeared to be a F_1_ hybrid between *B. g. grandii* and *B. g. benazzii*, with *B. rossius* mitochondria. The third one (c on Figure 3) could be triploid, bearing approximately equal contributions from each of *B. g. grandii*, *B. g. benazzii* and *B. r. redtenbacheri*, with *B. g. grandii* mitochondria. All three hybrid females had very high relative heterozygosity (Figure 3D).

Ancestry analyses per scaffold revealed no evidence for large-scale genome replacement in hybridogens (as would be expected, for example, if a hybridogenetic individual would have transmitted a *B. grandii* instead of a *B. rossius* homolog, or if chromosomes occasionally recombined before genome elimination). By contrast, three individuals of the triploid, parthenogenetic *B. lynceorum* had ancestries corresponding to only two instead of the expected three species on the X chromosome (Figure 4, Figure S4). Two of them were seemingly missing the *B. g. grandii* X, while one was seemingly missing the *B. r. redtenbacheri* X. The ratio of depth at the different alleles on polymorphic X-linked sites indicated that these individuals were likely triploid on the X chromosome, suggesting that one homolog may have been replaced, rather than lost, during evolution (see Supplementary material, Figure S5).

**Figure 4:**
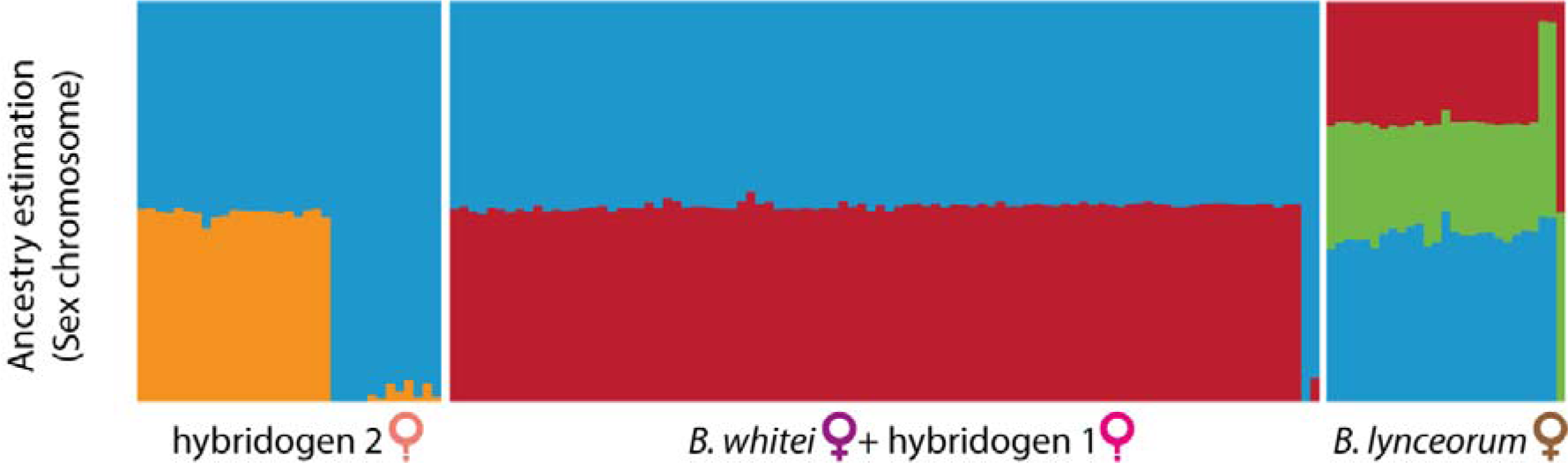
X chromosome ancestry in hybrids. Note that the (rarely produced) males of the hybridogenetic lineages feature, as expected, a pure *B. r. redtenbacheri* ancestry at their sex chromosome given the XX/XO sex determination. Note the ancestry corresponding to two instead of the expected three species on the X chromosomes of three *B. lynceorum* individuals.

Heterozygosity was lower than expected in all except one diploid hybrids (one individual of the hybridogen 1; Figure 5A, B). This lower heterozygosity was due to local heterozygosity drops that were spread throughout the genome and were mostly shared among individuals within lineages, and even mostly between the hybridogens 1 and *B. whitei* (Figure 5C, Figure S6).

**Figure 5:**
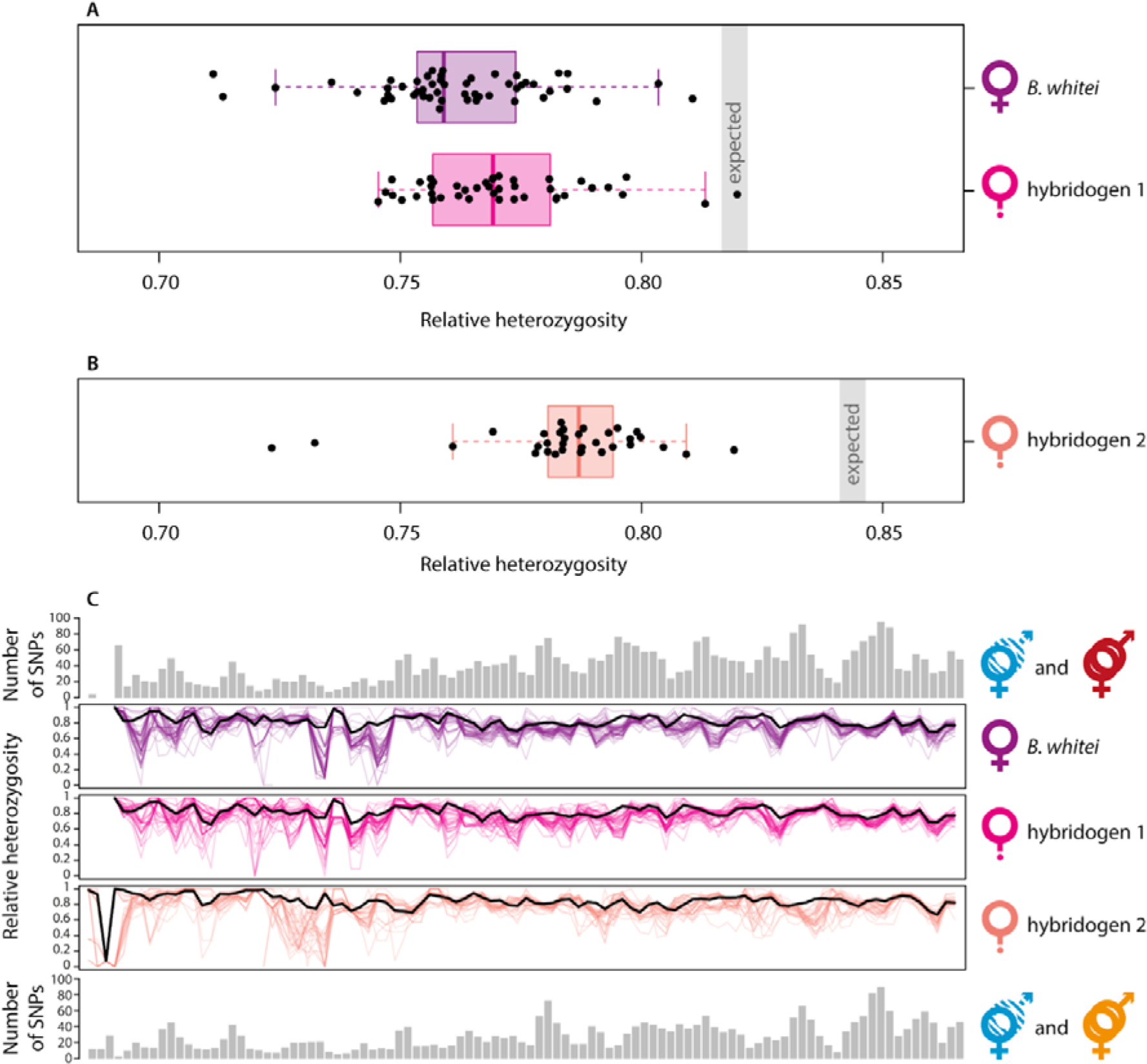
Observed relative heterozygosity on autosomes in hybrids compared to expected heterozygosity in newly-formed hybrids, computed based on allele frequencies in the sexual populations. A: in hybrids with *B. g. grandii* as paternal ancestor. **B**: in hybrids with *B. g. benazzii* as paternal ancestor. The boxplots show observed heterozygosity values and the grey areas are the range between minimum and maximum expected values (out of 1000 replicates). **C**: distribution of observed and expected heterozygosity along the first 50 Mb (100 sliding windows) of scaffold 1, which is representative of all autosomes. Top and bottom: number of SNPs per window genotyped in at least 3 individuals of both *B. r. redtenbacheri* and *B. g. grandii* (top) or *B. g. benazzii* (bottom). Middle panels: heterozygosity in each individual (colored lines), and expected heterozygosity based on the parental allele frequencies (averaged across 1000 replicates; thick black lines). A high-resolution version of this figure for the whole genome is available in Figure S6.

We found large variations in relative heterozygosity among the different lineages (Figure 5). Asexual populations of *B. r. redtenbacheri* had the lowest relative heterozygosity, followed by, in increasing order: sexual populations of *B. r. redtenbacheri*, *B. g. grandii* and *B. g. benazzii*, *B. atticus*, *B. whitei*, and *B. lynceorum*. Because of low sample size (7 individuals), heterozygosity in *B. g. maretimi* was not significantly different from any of the sexual species nor the asexual populations of *B. r. redtenbacheri*.

**Figure 5:**
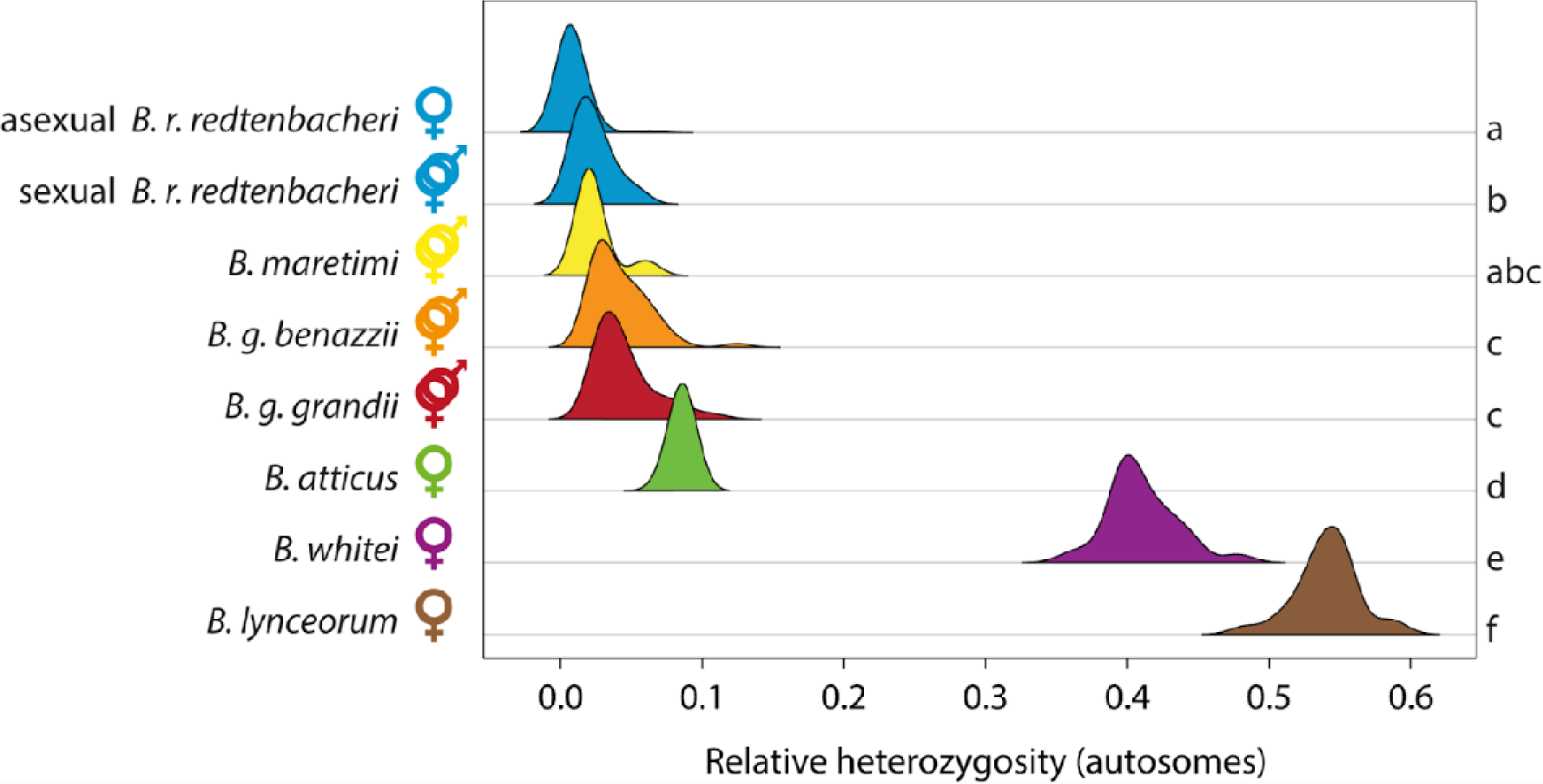
Distribution of relative heterozygosity on autosomes of the different sexual and parthenogenetic lineages of *Bacillus*. Different letters on the right indicate significant differences. Colors and symbols as in Figure 2.

## Discussion

We revisited the *Bacillus* species and lineages from Sicily, using RAD sequencing and COII barcoding for 396 wild-caught individuals, and corroborated the original species descriptions and genomic composition of hybrid species and lineages with alternative reproductive modes. We found that chromosome 2 was the sex chromosome in all species. Our chromosome-level reference genome of *B. rossius redtenbacheri*, the maternal ancestor of all hybrids, then allowed us to study ancestry variation along chromosomes. All 75 individuals from the two hybridogenetic lineages had an F_1_ hybrid ancestry across their entire genome, retaining most of their heterozygosity. This means that hybridogenesis was never leaky, i.e., elimination of the *B. grandii* genome was always complete, and the two genomes never recombined, consistent with the elimination of the *grandii* genome prior to meiosis (Marescalchi & Scali, 2001). Absence of recombination was also shown by genomic *in situ* hybridization in hybridogenetic *Pelophylax* frogs (Zaleśna et al., 2011) and *Hypseleotris* fish (Majtánová et al., 2021), and by whole-genome resequencing in the basidiomycete fungus *Cryptococcus neoformans* (Yadav et al., 2021). This consistency across very different hybridogenetic taxa suggests that complete lack of recombination may be required for triggering hybridogenesis in hybrid individuals and/or be key for the long-term persistence of hybridogenetic taxa.

We also found a complete maintenance of F_1_ hybrid ancestry across the entire genome for all 52 individuals of the obligate parthenogen *B. whitei*. This is consistent with the described mechanism of parthenogenesis for this species, endoduplication, whereby chromosomes undergo an extra duplication prior to meiosis and the duplicates then segregate during meiosis I, thus leading to genetically identical offspring that retain the hybrid constitution of their mother (Marescalchi & Scali, 2001). The same was found in 22 out of the 25 triploid *B. lynceorum*. The three remaining individuals bore X chromosomes from two instead of the three expected parental species. The fact that they were triploid for the X chromosome indicates that they carry two copies of the X chromosome from one of their parental species. This would be in line with a dysfunctional endoduplication, as endoduplication is also the described mechanism of parthenogenesis in this species (Marescalchi et al., 1991). If pairing and segregation occasionally occurred between homologs from different species rather than chromosome duplicates, this could lead to two chromosomes from the same parental species ending up in the oocyte. Such pairing between homologs rather than duplicates is particularly likely for the X chromosome, given the strong synteny and slow evolution of the 450 My old insect X chromosome (Li et al., 2022; Parker et al., 2022; Toups & Vicoso, 2023).

While hybrid lineages overall retained their hybrid constitution, we found that heterozygosity was slightly lower in the two hybridogenetic lineages as well as in the parthenogenetic *B. whitei* than expected from the divergence between the sexual populations of *B. grandii* and *B. r. redtenbacheri*. Loss of heterozygosity was also observed on rare occasions at some allozyme loci in *B. whitei* and *B. lynceorum* (Mantovani et al., 1992). Similar patterns are found in some, but not all, asexual hybrid animals. In the Amazon molly *Poecilia formosa*, heterozygosity is slightly reduced as a consequence of gene conversion, which is biased towards one parental species (Tiedemann et al., 2005; Warren et al., 2018). The same was found in the cricket *Warramaba virgo* (Kearney et al., 2022). Gene conversion also appears to reduce heterozygosity in asexual lineages of the water fleas *Daphnia pulex* and *D. obtusa* (Xu et al., 2011), although their hybrid origin is not clear (Tucker et al., 2013). Larger heterozygosity losses occur via occasional recombination in salamanders of the genus *Ambystoma* (Bi & Bogart, 2006). On the other hand, no net losses were observed in *Cobitis* loaches (Janko et al., 2021), likely because the effect of gene conversion is compensated by the increased heterozygosity resulting from the independent evolution of the parental genomes, a pattern also reported for the fissiparous ribbon worm *Lineaus pseudolacteus* (Ament-Velásquez et al., 2016).

Two biological and one technical hypotheses could explain why hybrids have lower heterozygosity than expected. First, gene conversion can cause local drops in heterozygosity. Formally testing for gene conversion would require a marker density higher than the typical tract length of gene conversion (a few hundred bases; Casola et al., 2010; Setter et al., 2022). This was not possible with our RAD data, which provided only a few dozens of markers per Megabase). Second, the two parental genomes may have diverged more quickly in their respective sexual species than in the hybrids, owing for example to differences in substitution rates. This is likely, as the *rossius* genome in hybridogens and both genomes in *B. whitei* were only affected by mutations in the female germline, which are typically much lower than male germline mutation rate (Bergeron et al., 2023), while the genomes of the sexual species are affected by mutations both in the male and female germlines. Finally, allelic dropout could contribute to the lower observed heterozygosity than expected. Most (> 80%) loci showed fixed differences between the two parental species, meaning that all individuals of the parental species are homozygous, and allelic dropout only occurs at heterozygous sites. On the other hand, heterozygous sites can be affected by allelic dropout in hybrids, which will reduce observed heterozygosity. For this reason, conducting the same analysis for *B. lynceorum* was not possible as this species is triploid, which results in even lower sequencing depth per allele, and thus higher allelic dropout. Investigating the genomic distribution of heterozygosity in *B. lynceorum* and the impact of dropout on observed heterozygosity in diploid hybrids would require more extensive genomic data and is a task for future research.

Independently of the explanation for the reduction of observed as compared to expected heterozygosity in hybrids, the variation of heterozygosity levels among the three parthenogenetic species (*B. whitei, B. lynceorum, B. atticus*) and the parthenogenetic populations of *B. rossius* were consistent with the different proximate mechanisms underlying parthenogenesis in each case. The lowest heterozygosity level was found in parthenogenetic populations of *B. r. redtenbacheri*. Pijnacker (1969) found that females of this species achieve parthenogenesis via gamete duplication, which generates completely homozygous individuals. Haploid eggs are produced via regular meiosis. In some of the egg nuclei, the chromatids replicate but do not segregate, which results in a diploid, but fully homozygous embryo. The low levels of heterozygosity that we observed in these populations are likely an artifact caused by structural variants, as has been observed elsewhere (Jaegle et al., 2023; Jaron et al., 2022; Larose et al., 2023). In *B. atticus*, heterozygosity was higher than in all sexual species, consistent with the hypothesis that this species might be of hybrid origin, but involving currently unknown species or populations (Bullini & Nascetti, 1989). The proposed mechanism of parthenogenesis in this species is central fusion automixis (Marescalchi et al., 1993), whereby the oocyte fuses with one of the products of the first meiotic division (i.e. homologous chromosomes rather than chromatids which are separated at the second meiotic division). Heterozygosity is thus conserved in all regions that have not recombined (Suomalainen et al., 1987). Most heterozygosity is still expected to be lost over the long term, except in regions that never recombine (e.g. near the centromeres or, in hybrids, in regions that are too divergent to recombine) or where heterozygosity is maintained via selection. A reference genome assembly for *B. atticus* would be needed to locate the retained heterozygosity in the genome of this species, as potential chromosome rearrangements and small-scale translocations during the independent evolution of *B. atticus* and *B. r. redtenbacheri* (our reference genome) prevent us from doing so with the current data. Finally, heterozygosity was even higher in *B. whitei* (in contrast with previous findings from a few allozyme markers that heterozygosity was higher in *B. atticus* than in *B. whitei*; Nascetti & Bullini, 1980), and even more so in the triploid *B. lynceorum*. This is consistent with the hybrid composition of *B. whitei* and *B. lynceorum* and with the fact that they achieve parthenogenesis via endoduplication (as discussed above), thus leading to the complete maintenance of heterozygosity (Marescalchi et al., 1991). The fact that the facultative parthenogenetic species (*B. rossius*) is non-hybrid and loses heterozygosity while obligate asexuals are hybrid and retain high levels of heterozygosity parallels what is found in whiptail lizards (Lutes et al., 2010; Barley et al., 2022; Ho et al., 2023). More generally, the higher retention of heterozygosity in hybrid vs non-hybrid asexuals appears to be a general pattern across taxa, regardless of the proximal mechanisms of parthenogenesis (Jaron et al., 2022).

The fact that the genus *Bacillus* encompasses the full breadth of reproductive modes found in animals raises the question of whether they are predisposed to unusual reproduction. Facultative parthenogenesis appears to be widespread in stick insects and often (but not always) occurs via gamete duplication (White, 1973; Larose et al., 2023), suggesting that it might be ancestral to Phasmatodea. Why none of the subspecies of *B. grandii* are able to reproduce by parthenogenesis is yet unclear. All populations of *B. grandii* appear to be small and isolated, which could result in a high load of recessive deleterious mutations. Under gamete duplication, these mutations would be exposed and their cumulative fitness effects could be lethal. The absence of parthenogenesis in *B. grandii* could thus be caused by population genetic processes rather than mechanistic constraints. Whatever the explanation for the lack of parthenogenesis in *B. grandii*, facultative parthenogenesis is unlikely to developmentally predispose *Bacillus* for other unusual reproductive modes. Indeed, hybridization between different species led to two different proximate parthenogenesis mechanisms (central fusion automixis in *B. atticus* and endoduplication in *B. whitei* and *B. lynceorum*). In the case where the repeated evolution of alternative reproductive modes in *Bacillus* would be due to a developmental predisposition, one would expect a conserved mechanism. Finally, the diversity of reproductive modes displayed by lineages with contributions from *B. rossius* and *B. grandii* raises the question whether there are specific features in their respective genomes that lead to unusual reproduction. It may be that their divergence upon hybridization was within the specific range of differentiation where hybridization was still possible, but the amount of divergence was enough to disrupt meiosis and associated reproductive mechanisms, as predicted by the balance hypothesis (Moritz et al., 1987) and observed in whiptail lizards (Barley et al., 2022) and loaches (Marta et al. 2023). Understanding whether the diversity of reproductive modes in *Bacillus* hybrids is the consequence of repeated hybridization or secondary diversification is thus an exciting task for future research.

In addition to previously described hybrid lineages, we found three new hybrids, which may represent new lineages with alternative reproductive modes. The first one was a female, seemingly F_1_ between the parthenogen *B. atticus* and *B. g. grandii*. The two species live in sympatry in the area where this individual was collected. The fact that it derived its mitochondria from *B. atticus* implies that this species was its maternal ancestor and that *B. atticus* females have retained the capacity to reproduce sexually. Taken together with the documented production of rare males in this species (Scali, 2013), this opens the possibility for cryptic sex in this otherwise obligate parthenogenetic species, which could contribute to its long-term maintenance (as for example in some species of *Timema* stick insects; Freitas et al., 2023; Larose et al., 2023).

The second new hybrid was a female with an apparent F_1_ composition between *B. g. benazzii* and *B. g. grandii*, and with *B. rossius* mitochondria. The third one was a female with an approximately equal contribution from each of *B. g. benazzii, B. g. grandii*, and *B. r. redtenbacheri*, with *B. g. grandii* mitochondria. Hybridization between the two subspecies of *B. grandii* is puzzling, as they were thought to be distributed on opposite sides of Sicily, about 300 km away from each other (Scali et al., 1995). The high heterozygosity of these hybrids comparable to F_1_ individuals points to a recent origin, which would suggest that a relictual population of *B. g. benazzii* might exist in the South-East of the island. High heterozygosity could also be maintained over generations via (functionally) apomictic parthenogenesis. Interestingly, the existence of these hybrids confirms that hybridization can still occur between the different *Bacillus* species (see Mantovani et al., 1996). However, our results indicate that this did not lead to gene flow between the parental species.

In conclusion, the results of our extensive RADseq and COII genotyping of field-collected *Bacillus* stick insects confirm older genetic descriptions of *Bacillus* hybrids and their parental species, as well as their ancestral genomic compositions. While the previous analyses were restricted to a few loci, our new chromosome-level reference genome of *B. r. redtenbacheri* enables us to infer that all ancestry estimates are representative of the whole genome. In addition, we discovered three new types of hybrids, which might represent lineages that were so far overlooked. We found that the two hybridogenetic lineages have an F_1_ constitution along the whole genome, showing no evidence for “leaky” hybridogenesis. By contrast, some triploid asexual hybrids have replaced one X chromosome inherited from one parental species with a copy from another parental species. Overall, our genetic identification and characterization of field-caught individuals paves the way for experiments to illuminate the proximate molecular mechanisms of genome elimination under hybridogenesis and for studies on the origin and diversification of reproductive modes in the genus. Our results and the genomic resources presented here will serve as a tool for using *Bacillus* as a new model organism in the future.

