## Supplementary text and figures for "Evolution of alternative reproductive systems in *Bacillus* stick insects"

### Supplementary Material

#### Sex chromosome identification

Sex determination follows an XX/X0 system, where females have two copies of the X and males have only one in both *B. rossius* (Scali & Mantovani, 1989) and *B. grandii* (Marescalchi and Scali 1990). We identified which of our 18 scaffolds was the sex chromosome by looking at male / female sequencing depth differences. Males are expected to have half the depth of females on the X chromosome, and similar depths on other chromosomes. We first standardized depth per individual by dividing the depth at each locus by the individual’s average depth across all loci. We then calculated male / female depth ratio at each SNP as the average standardized depth in males, divided by that of females. We did it separately for *B. r. redtenbacheri* (57 females, 10 males), *B. g. grandii* (24 females, 29 males) and *B. g. benazzii* (28 females, 11 males). We found that male / female depth ratio was much lower on the whole of scaffold 2 than on all other scaffolds, in all (sub-)species (Figure S1). This indicates that scaffold 2 in our assembly is the sex chromosome, which is conserved in *Bacillus*. We further verified whether our findings were consistent with previous work by locating the *Mdh-2* gene, which was suggested to be X-linked (as males were invariably homozygous and females occasionally heterozygous; Tinti and Scali, 1995). We blasted the *Mdh-2* protein of *Drosophila melanogaster* (GenBank Accession Number NT_033777.3, bases 18226396 – 18228718) on our assembly using tblastx v2.12.0+. The ten best hits (all with e-values < 2*10^-27^) were located on the same region of scaffold 2, between positions 124’292’616 and 124’321’452. No other hits had an e-value < 1. This confirms that *Mdh-2* is X-linked in *B. rossius*.

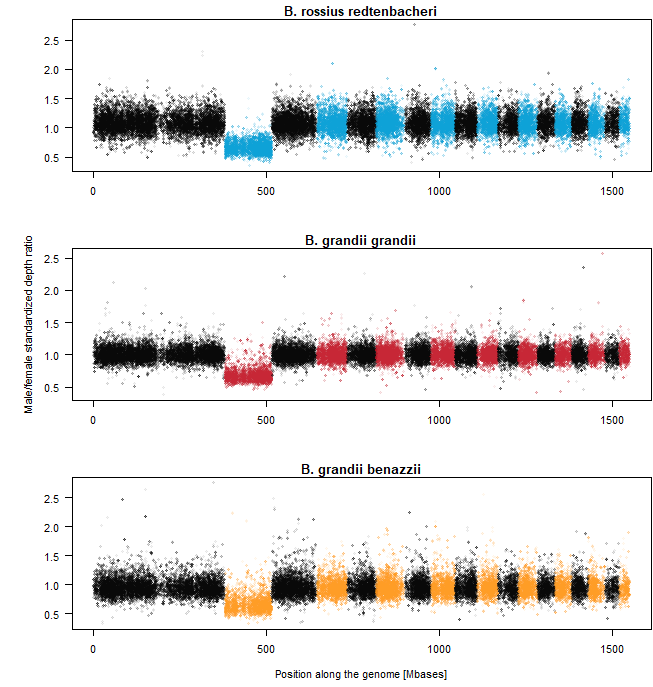

**Figure S1**: Manhattan plot of male / female standardized depth ratio for all three *Bacillus* subspecies where enough males and females were available (top: *B. r. redtenbacheri*; middle: *B. g. grandii*; bottom: *B. g. benazzii*). Each point is one SNP. Alternated colors designate different chromosomes.

#### Genotypic sex of individuals

We used standardized depth ratio at the X chromosome vs autosomes to sex field-collected individuals for which the phenotypic sex was unknown (because they were collected as juveniles). Unlike in the previous section, the depth ratio in this case was computed for each individual separately, averaging across X-linked and autosomal loci respectively. We found that X / autosomes depth ratios were centered a bit higher than the expected 0.5 in males (because males are X0 and have one copy of the X, while females are XX and have two) and 1 in females (Figure S1, Figure S2). We hypothesize that this may be due to the accumulation of structural variants on the X, which results in the X being enriched in paralogs. However, the distribution of male and female X / autosome depth ratios were not overlapping: all males had a depth ratio inferior to 0.72, while all females had a depth ratio higher than 0.9 (Figure S2). Individuals of unknown sex with a depth ratio inferior to 0.72 were thus considered males, and those with a depth ratio higher than 0.9 were considered females.

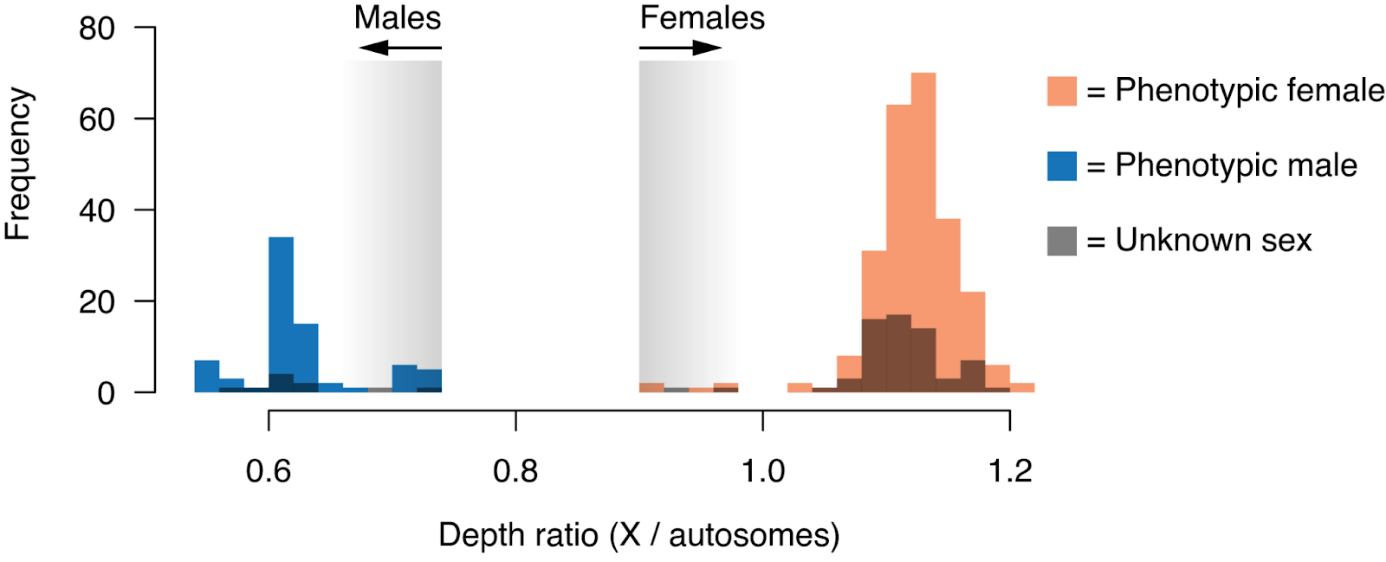

**Figure S2**: Distribution of standardized X / autosome depth ratio in phenotypic males and females, and individuals of unknown phenotypic sex.

**Figure S3** (next page): Maximum-Likelihood phylogeny of 695 bp of the mitochondrial gene COII. Colored arrows represent reference sequences from Mantovani et al. (2001), colored according to their species following the color code of Figure 1. Dark blue = *B. rossius* subspecies not present in Sicily. The colored rectangles define mtDNA haplotypes, which are used in Figure 2A,B.

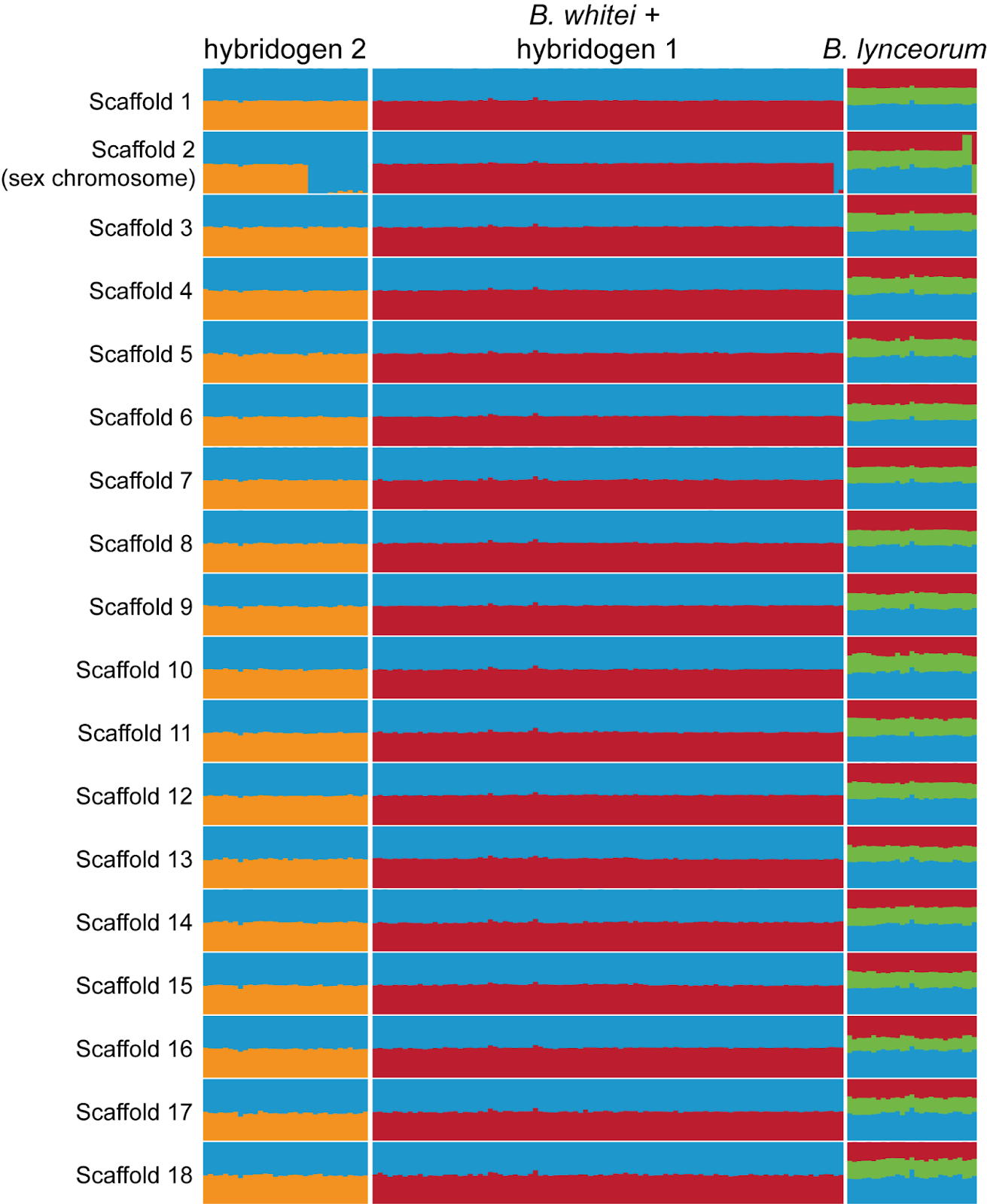

**Figure S4**: Ancestry proportion per chromosome for the three major types of hybrids. The hybridogens with pure *B. rossius* ancestry at their sex chromosome are males. Note the ancestry of the X chromosome in three *B. lynceorum* individuals, indicating contribution from two instead of the expected three parental species.

#### Loss or replacement of an X chromosome in some *B. lynceorum*?

Three out of 25 *B. lynceorum*, which are triploid hybrids with one complete haplome of each of three different parental species (see Figure 2B main text), had X chromosome assignments corresponding to only two instead of three parental species (Figure 4, main text; Figure S4). Two hypotheses could account for this observation: (i) they have lost an X chromosome and are therefore diploid for the X, or (ii) they are still triploid but the copy derived from one of the parental species was replaced by a second one from one of the other parental species. In order to distinguish between these two hypotheses, we looked at allelic depth ratio at heterozygous sites. We define allelic depth ratio as the proportion of reads corresponding to the allele with the highest coverage divided by the total number of reads at each locus with heterozygous genotype. Allelic depth ratios at heterozygous, biallelic SNPs are expected to be centered around 0.5 in diploids and 0.67 in triploids. We plotted the distribution of allelic depth ratio for the three focal individuals, along with ten randomly selected *B. lynceorum* and ten randomly selected *B. whitei*, used as null expectations for triploids and diploids, respectively. We found that the three *B. lynceorum* with X chromosome assignments corresponding to two species had allelic depth ratios on the X chromosomes similar to other triploid *B. lynceorum* and different from diploid *B. whitei* (Figure S5). This indicates that they have not lost an X chromosome; rather, they seem to have replaced one of their X homologs with a second copy from an X homolog from another parental species.

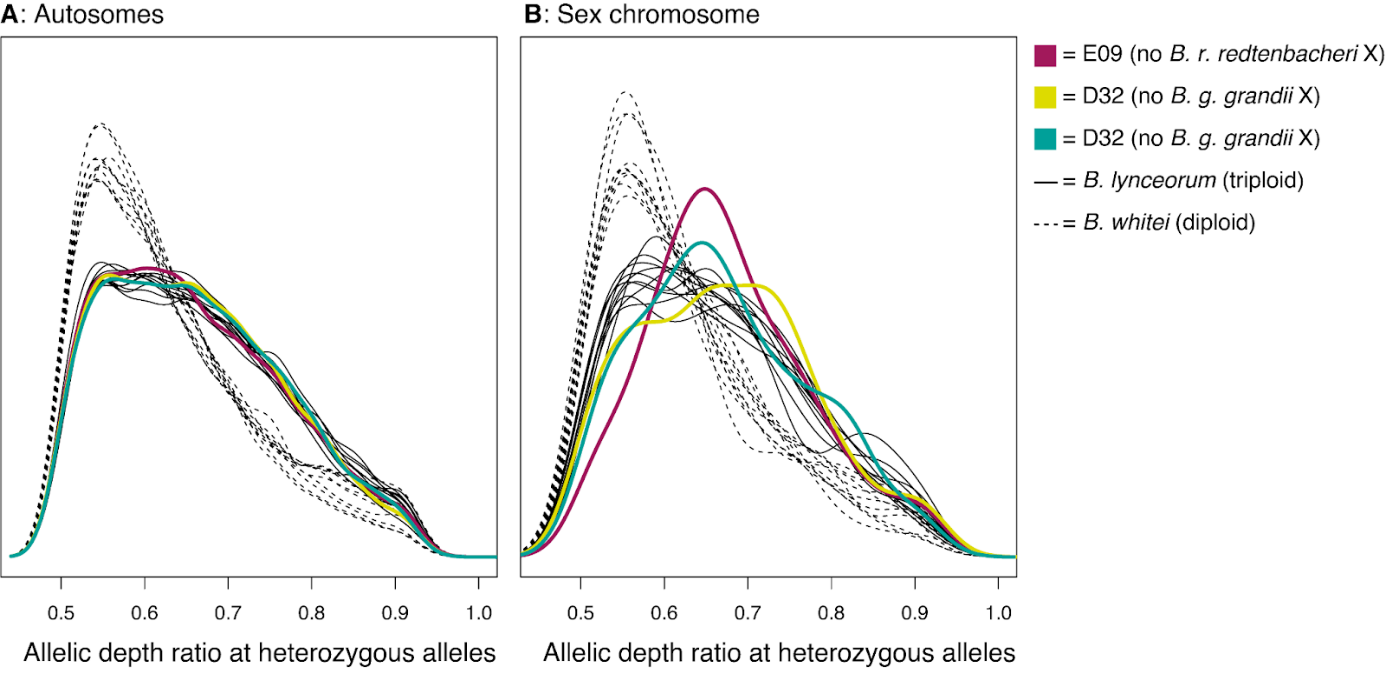

**Figure S5**: Distribution of allelic depth ratios at heterozygous sites for (A) autosomes and (B) the X chromosome, in ten randomly selected diploid *B. whitei* (dashed lines), ten randomly selected triploid *B. lynceorum* (solid lines), and the three *B. lynceorum* with X chromosome assignments corresponding to only two species (thick colored lines).

**Figure S6** (separate file): Observed and expected heterozygosity along the whole genome for the three lineages of diploid hybrids. See Figure 4 (main text) for detailed caption.

#### Ploidy level of the hybrid between B. atticus and B. g. grandii

We used the same allelic depth ratio approach as above to assess the ploidy of individual D34 (labelled as “a” in Figure 2), the hybrid between *B. atticus* and *B. g. grandii*. We used all 13 individuals of *B. atticus* as references (since heterozygosity is higher in *B. atticus* than in *B. grandii*) We found that the allelic depth ratio distribution of individual D34 was centered around 0.5 and not 0.67, identical to that of other *B. atticus*, indicating that it is diploid (Figure S7).

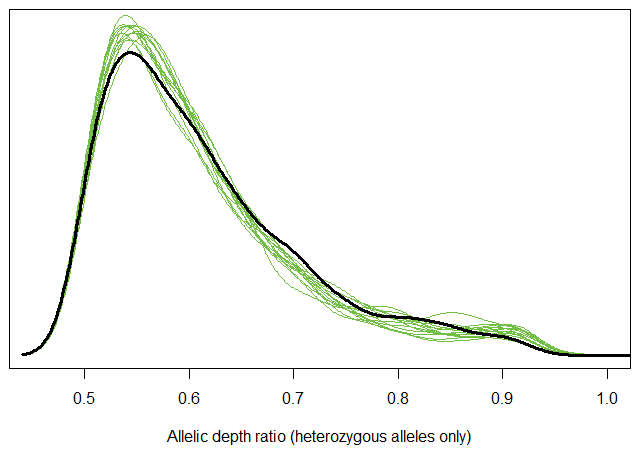

Figure S7: Allelic depth ratio distribution at heterozygous sites (genome-wide) for B. atticus individuals (thin green lines) and individual D34, the hybrid between *B. atticus* and *B. g. grandii* (thick black line).
