## Supplementary figures and images for "Evolution of alternative reproductive systems in *Bacillus* stick insects"

### Supplementary Figure 3

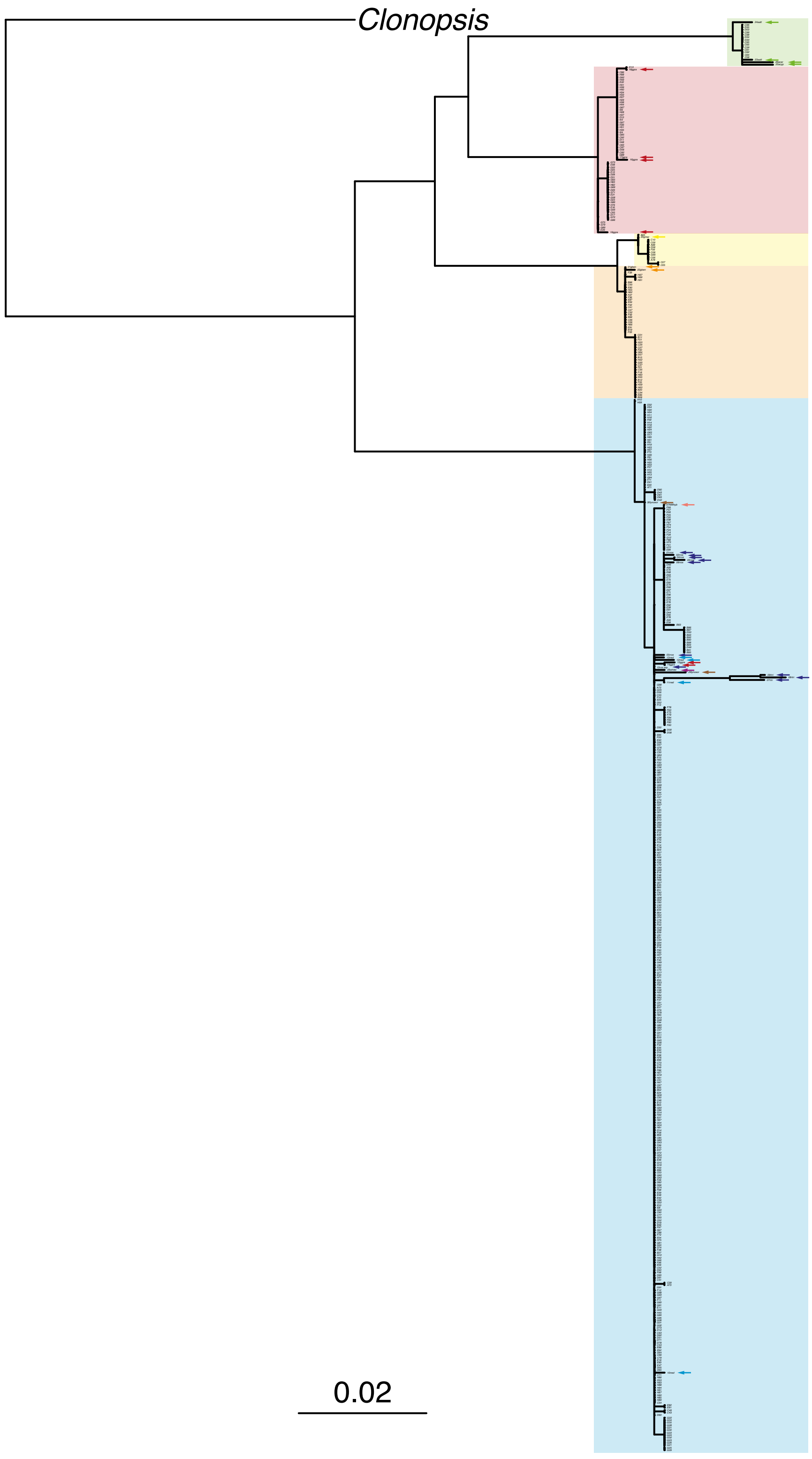

### Supplementary Figure 6

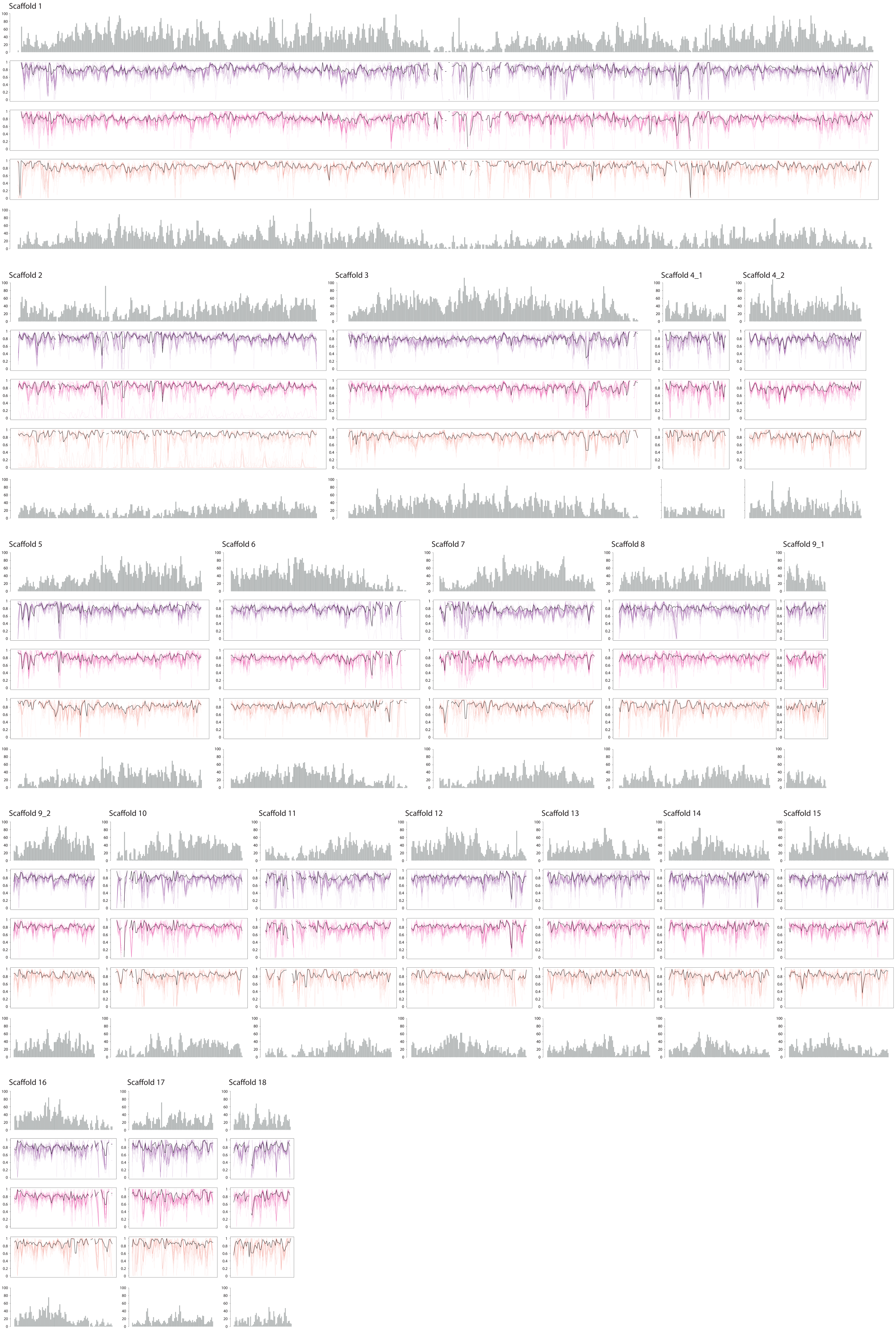
